# Intraguild parasitism promotes the persistence of facultative hyperparasitoids by extending temporal host availability

**DOI:** 10.1101/2025.02.25.640037

**Authors:** Hua Wang, Nuo Xu, Chi Xu, Jinbao Liao, Shaopeng Wang, Shucun Sun, Xinqiang Xi

**Author notes:** Hua Wang and Nuo Xu contribute equally to this study.

## Abstract

Intraguild predation (IGP) is pervasive in natural food webs, although theoretical models predict restricted parameter space for the coexistence of IG-prey and IG-predators. One potential mechanism is that IGP persistence might be increased when IG-prey extend resource availability for IG-predators. Here we test this hypothesis using manipulative experiments in combination with mathematical modeling. Our experiment system includes *Drosophila* flies (*Drosophila melanogaster*, basal resources), primary larval parasitoids (*Asobara leveri*, IG-prey) and facultative pupal hyperparasitoid (*Pachycrepoideus vindemiae*, IG-predator). We measured the performance of facultative pupal parasitoids with and without the primary larval parasitoid. Our study revealed that larval parasitoids significantly increased the number of pupal parasitoids emerged from limited host resources, due to the extended resource availability to facultative hyperparasitoid. Specifically, immature primary parasitoids within *Drosophila* pupae can be explored by facultative hyperparasitoids for up to 14 days, while the healthy fly pupae are available to hyperparasitoids for only 5 days. Despite a significant reduction in offspring fitness of the facultative hyperparasitoids when reared on larval parasitoids versus healthy *Drosophila* pupae, adult female facultative hyperparasitoids showed no significant preference for healthy fly pupae over immature primary parasitoids. Our theoretical models demonstrated that an extended development period of the primary parasitoid can promote the persistence of the facultative hyperparasitoids and allow their coexistence with the primary parasitoid, although too prolonged development could drive the extinction of the primary parasitoid. As IG-prey often take longer time to develop after consuming the shared resources, extension of temporal resource availability could be a general mechanism contribute to the persistence of intraguild predators.

**Open research statement:** Data are not yet available, but all of the original experimental data has been displayed in figures 2∼4. The data will be made publicly available upon manuscript acceptance and archived in the Dryad Digital Repository.

## Introduction

Competition and predation are two primary forces that determine the persistence of species and the structure of biological communities (Chesson & Kuang, 2008). Although these two processes are often studied separately, species competing for shared resources may also engage in trophic interactions. Intraguild predation (IGP), depicting the predation between consumers competing for shared resource (Polis & Myers, 1989), has been observed in both free-living predators and parasitic consumers (Borer & Briggs, 2007). Theoretical models of IGP modules predicted that the interplay between predation and exploitative competition made it difficult for the IG-predators and IG-prey to coexist in food webs (McCann & Hastings, 1997). However, detailed observation and application of molecular techniques have revealed that IGP are ubiquitous across diverse ecosystems (Arim & Marquet, 2004; Rosenheim, 2007). Therefore, increasing efforts are devoted to illustrate mechanisms contributing to the stability of IGP (Liao et al., 2017; Liao & Bearup, 2020) by incorporating detailed life-history traits of consumers (Sentis et al., 2014; Gutgesell et al., 2022; Bassar et al., 2023).

Theoretical models predict that the stability of IGP modules requires a balance between the consumptive and explorative interactions between IG-prey and IG-predators, which can be dependent of resource supply (Holt & Polis, 1997). At intermediate-level resource productivity, IGP promotes species coexistence when the IG-prey is a superior competitor compared with the IG-predator (Holt & Polis, 1997; Borer et al., 2003). At the high and low ends of the productivity gradient, however, IG-prey and IG-predators could be excluded through apparent and exploitative competition, respectively (Polis & Myers,1989; Borer et al., 2003). Previous studies have revealed several mechanisms that enable IG-prey to counteract the disadvantage of being preyed upon (Rosenheim et al., 1995; Vance-Chalcraft et al., 2007). For instance, invulnerable development stages can decrease the mortality of IG-prey (Borer, 2002), and extraguild prey may also alleviate the predation pressure on IG-prey (Lucas & Coderre, 1998). Conversely, how IG-predators survive the intense competition with IG-prey at low productivity remains less clear (Holt & Polis, 1997; Morin, 1999).

Theoretical studies have revealed at least two ways that could increase the persistence of top predators amidst extensive resource competition. Firstly, adaptive foraging by IG-predators may increase the proportion of IG-prey consumption upon basal resources at low productivity (Murdoch & Oaten, 1975; May, 1977; Gutgesell et al., 2022), alleviating resource competition for shared resources and enabling the rapid recovery of shared resources (Holt & Polis, 1997; Navarrete & Menge, 2000; Moeller & Neubert, 2019). To ensure stable coexistence of consumers, IG-predators need to have lower preference and feeding efficiency on the IG-prey than on the basal resources (McCann & Hastings, 1997; Amarasekare, 2007). Therefore, more nutrient of IG-predators come from basal resources rather than IG-prey (Polis & Myers, 1989; Wang & Brose, 2019). Omnivory models are constructed instead to explore how IG-prey survive predation when they are preferred over basal resources (McCann & Hastings, 1997). Although IG-prey are generally less vulnerable than basal resources (Müller & Brodeur, 2002; Zheng et al., 2021), IG-predators may obtain a higher proportion of nutrients from IG-prey than shared resources at low productivity (Kratina et al., 2012). For example, omnivorous amphipods periodically transit their diet from predominantly consuming phytoplankton (shared resource) in the open water at warm season to predominantly feeding on zooplankton (IG-prey) under ice cover during the winter (McMeans et al., 2015).

Secondly, while IG-prey are typically perceived as depleting shared resources, they can paradoxically enhance resource availability for the IG-predator through non-consumptive mechanisms (Banerji & Morin, 2014). For instance, rabbits may reduce their time outside shrubs in the presence of swift foxes (IG-prey), thereby increasing their exposure to wolves (IG-predators) at night (Thompson & Gese, 2007). Recent research indicates that IG-prey may facilitate the transmission of viruses to basal resources, subsequently increase the vulnerability of resources to top predators (Flick & Coudron, 2020). In addition, IG-prey can enhance the quality of resources for IG-predators. For example, mouth width of *Tetrahymena vorax* (IG-predator) increase considerably when they consuming protozoa such as *Colpidium kleini* (IG-prey), enabling them to prey on larger prey (Banerji & Morin, 2014). Furthermore, IG-prey generally have longer lifespans compared to basal resource species, consumption of basal resources by IG-prey can extend the temporal energy available to top predators, particularly when these basal resource species are nearing the end of their life cycle (Gutgesell et al., 2022). For example, caribou can store the energy fixed by short-lived annual grasses and transfer it to omnivorous bears over a longer period (Rayl et al., 2018).

Parasitoids are dominant natural enemies of invertebrate herbivores in many ecosystems, which often utilize a narrow range of host species at specific developmental stages (Godfray, 1994). Variations in the efficiency of exploit resources at different trophic levels are particularly significant for facultative hyperparasitoids whose fitness is largely determined by the encounter rate with their hosts (Lafferty et al., 2008). While the dynamics of parasitoids have primarily been studied within simplified modules of trophic interactions, such as host-parasitoid or host-parasitoid-hyperparasitoid systems (Hassell, 2000), certain (hyper)parasitoid species may engage in interactions resembling IGP between free-living predators (Borer & Briggs, 2007). In particular, IGP could emerge when multiple parasitoids oviposit successively during the development of the same host. Parasitoids that oviposit in larval hosts are often consumed by subsequent parasitoids that can attack both healthy hosts in their later developmental stages and hosts that have been parasitized earlier (Borer, 2002; Borer & Briggs, 2007; Frago, 2016). Furthermore, due to the limited ability to identify whether hosts have been parasitized by primary parasitoids (IG-prey), facultative hyperparasitoids (IG-predators) often exploit different hosts in a frequency-dependent manner (Sieber & Hilker, 2011). Although primary parasitoids suppress the population growth of host (Borer, 2002), they can increase the host quality and exposure period to facultative hyperparasitoids. For instance, herbivores parasitized by koinobiont primary parasitoids at the larval stage may be stimulated to ingest more nutrients, resulting in larger pupae that provide more nutrients to facultative pupal hyperparasitoids (Jervis & Kidd, 1986; Xi et al., 2015).

*Drosophila* flies are one of the most diverse insects in the wild, serving as hosts for more than 100 parasitoid species during their larval and pupal stages (Lue et al., 2021). In this study, we experimentally investigated how the larval parasitoid, *Asobara leveri*, alters host availability to the facultative pupal parasitoid *Pachycrepoideus vindemiae*. Given that (a) the fecundity of *A. leveri* is significantly higher than that of *P. vindemiae*, and (b) the period during which *P. vindemiae* can utilize hosts already parasitized by *A. leveri* is significantly extended, we hypothesize that IGP will increase the persistence of facultative hyperparasitoids by extending temporal host availability, and increase the productivity of two parasitoid species coexisting under the specific developmental time of *A. leveri*. To illustrate this, we constructed a stage-structured IGP model to examine how the extended temporal availability of hosts by larval parasitoids affects the persistence of pupal parasitoid.

## 1 Materials and Methods

### 1.1 Insect life histories and rearing

*A. leveri* is a koinobiont solitary parasitoid species that oviposits in the 1∼4 days old *Drosophila* fly larvae. The parasitized fly larvae do not die until they develop into pupal stage. In comparison, *P. vindemiae* is a solitary facultative hyperparasitoid, with adult female *P. vindemiae* ovipositing in either healthy *Drosophlia* pupae or pupae that have already been parasitized by *A. leveri* at the larval stage.

Laboratory colonies of flies and wasps were collected from Zijinshan National Forest Park, Jiangsu Province, China, in 2023 and reared in insect rearing chamber at 25 ± 1 with a light: dark cycle of 16:8 hours and relative humidity of 60∼70%. *Drosophila* flies were cultured using a cornmeal-based artificial diet composed of yeast, agar, and sucrose. To obtain enough *D. melanogaster* for the experiments, we placed six plastic vials (diameter = 2.5 cm, height = 9.5 cm) with 5 ml artificial diet into a plastic box (length × height × width = 40 × 20 × 20 cm) covered with mesh. Approximately 40 pairs (♀:♂ = 1:1) of adult *Drosophila* flies were then released into the box to lay eggs in the vials for 24 hours. Once the fly eggs hatched into the first instar larvae, three pairs (♀:♂ = 1:1) of adult *A. leveri* were introduced into the vials to lay eggs for 24 hours. To obtain enough pupal parasitoids, six pairs of adult *P. vindemiae* were given access to 40 fresh pupae to lay eggs for 24 hours. The parasitoid offspring were fed with 10% sucrose water solution for use in experiments.

### 1.2 Effects of larval parasitoid on availability of host to pupal hyperparasitoid

To determine whether *A. leveri* increase the availability of hosts for *P. vindemiae* at different host density, we released 6, 12, and 25 pairs (♀:♂ = 1:1) of adult *Drosophila* flies into vials to lay eggs for 24 hours. An average of 60, 300 and 500 *Drosophila* fly larvae and pupae were observed in the three adult density treatments respectively, which would result in a host shortage in the low-density treatment and host redundancy in the high-density treatment. After *Drosophila* fly eggs developed into 2-day-old larvae, one pair (♀:♂ = 1:1) of *A. leveri* was introduced into the vials to lay eggs until they died. Once the *Drosophila* fly larvae pupated, one pair of *P. vindemiae* was introduced into each vial until they died.

We repeated the procedure to explore how larval parasitoids change the population density of pupal parasitoids by not introducing *A. leveri* at fly larval stage. Parasitoid wasps were fed 10% sucrose water solution. A total of 20 replicates were constructed for treatments with and without larval parasitoids respectively. The number of wasps that emerged from host pupae at different host densities were recorded. To investigate whether the increase in the number of *P. vindemiae* offspring is attributed to the availability of *A. leveri*, we dissected host pupae at intermediate densities, both with and without *A. leveri*. Predation of *A. leveri* by *P. vindemiae* was confirmed when larvae of both parasitoid species were observed within a single pupa.

To determine whether the larval *A. leveri* within *Drosophila* pupae triggers a longer availability period for *P. vindemiae* compared to healthy *Drosophila* pupae, we released 8 pairs (♀:♂ = 1:1) of adult flies into vials to lay eggs for 6 hours. After the eggs developed into 2-day-old larvae, one pair (♀:♂ = 1:1) of *A. leveri* was introduced into the vials to lay eggs for 24 hours. We then selected the *A. leveri*-parasitized pupae based on their darker puparium and reared them separately. Healthy pupae without larval parasitoids were conducted concurrently to determine the duration of availability for *P. vindemiae*. The time taken for fruit flies to develop from pupae to adults, as well as the time taken for *A. leveri* to develop from eggs to adults, was recorded. A total of 30 healthy or parasitized pupae were observed separately.

To examine the preference of *P. vindemiae* for healthy pupae and pupae with larval *A. leveri* at different developmental stages, we exposed 2 pairs of *P. vindemiae* to 10 healthy pupae together with 10 second instar *A. leveri* larvae (5 days old), pre-pupae *A. leveri* (10 days old) or pupal *A. leveri* (15 days old) for 24 hours. The healthy and *A. leveri*-parasitized pupae were randomly arranged on the vial walls to mitigate the influence of pupa position on *P. vindemiae* selection. All pupae were dissected under a stereomicroscope on the 5th day after introducing *P. vindemiae* to record the number of pupae parasitized. A total of 10 replicates were conducted for each of three developmental stages of *A. leveri*.

### 1.3 Fitness of *P. vindemiae* hosted by healthy fly pupae and larval *A. leveri*

To determine the fitness cost of *P. vindemiae* on pupae parasitized by *A. leveri*, we released 4 pairs (♀:♂ = 1:1) of adult *Drosophila* flies into vials to lay eggs for 24 hours. After the eggs developed into 2-day-old larvae, 3 pairs (♀:♂ = 1:1) of *A. leveri* were introduced into the vials to lay eggs for 24 hours. Once the larvae had pupated, pupae containing *A. leveri* of 5-, 10-, and 15-days old were selected for the experiment. Since lifespan and fecundity are closely positively related to the body size of *P. vindemiae* (Mariano-Macedo et al., 2020, Wang et al., 2021), and the body mass of *P. vindemiae* offspring emerging from pupae containing 10- and 15-days old *A. leveri* is similar, only pupae containing 5- and 15-days old *A. leveri* were used to measure the lifespan and fecundity of *P. vindemiae* offspring. 20 parasitized pupae were exposed to 2 pairs (♀:♂ = 1:1) of *P. vindemiae* for parasitism for 24 hours. We measured the emergence rate, sex ratio (proportions of male offspring), body mass, lifespan and fecundity of the progeny of *P. vindemiae*. A total of 15 replications were conducted for each developmental stage of *A. leveri*.

36 newly emerged female *P. vindemiae* from healthy and parasitized pupae were euthanized using ether, then dried at 65 °C for 5 days to weigh their dry mass. 20 female *P. vindemiae* were fed 10% sucrose water solution to measure their lifespan. To compare the fecundity of *P. vindemiae* offspring emerged from different hosts, 20 healthy pupae were exposed to a pair (♀:♂ = 1:1) of *P. vindemiae* to oviposit for 24 hours, during which the parasitoids were fed a 10% sucrose water solution. This procedure was repeated for each pair of parasitoids, transferring them into a vial with an equal number of fresh host pupae every 24 hours for 12 days. Typically, more than 65% of eggs are laid during the first 12 days post emergence. 10 females were used to measure the fecundity of *P. vindemiae* emerged from different hosts.

### 1.4 Data analysis

We employed generalized linear models (GLMs) with Poisson error distribution (link = “log”) to test if the number of *P. vindemiae* emerged significantly differed from healthy pupae and those with immature *A. leveri*, and if the number of hosts parasitized by *P. vindemiae* with *A. leveri* was significantly higher than that of healthy pupae without *A. leveri*. These GLMs were constructed using the ‘*anova.glm*’ function in the ‘*stats*’ package (Hastie & Pregibon, 2017). The difference in the length of time between healthy and parasitized pupae was compared using Student’s t-tests. The impact of *A*. *leveri* age on the survival rate of *P. vindemiae* was evaluated using GLMs with binomial error distribution (link = “logit”). Tukey post-hoc tests were performed using the ‘*glht*’ function in the ‘*multcomp*’ package (Bretz & Hothorn, 2016) whenever a significant difference was detected in GLMs.

Subsequently, we employed GLMMs in the ‘*lme4*’ package with Poisson error distribution to test whether *P. vindemiae* has a significant preference for healthy pupae over those parasitized by *A. leveri*. In the GLMM, individual parasitoids were treated as random effects, and then the significance of fixed effects (pupa type) was assessed using the ‘*Anova*’ function in the ‘*car*’ package. We employed Student’s t-tests to compare the body mass and sex ratio of *P. vindemiae* progeny emerging from healthy and *A*. *leveri*-parasitized pupae. A one-way analysis of variance (ANOVA) was used to test whether there were significant differences in the body mass and sex ratio of *P. vindemiae* progeny across different ages of *A. leveri*. We employed GLMs with Poisson error distribution to compare the fecundity and longevity of *P. vindemiae* progeny emerging from healthy and *A*. *leveri*-parasitized pupae. Tukey post-hoc tests were used to test whenever a significant difference in the fecundity and longevity of *P. vindemiae* progeny across different ages of *A. leveri*.

### 1.5 Theoretical model

As primary parasitoid tends to increase the exposure period of hosts to facultative hyperparasitoids in various parasitoid-host food web to varying extents (Godfray 1994), we construct a theoretical three-species IGP module based on an established stage-structure model (Borer & Briggs, 2007) to understand how parasitism-induced extension in the host exposure period (represented by a slower development of the primary parasitoid, i.e., lower *m_AL_*) influences the persistence of parasitoid species. Briefly, fly larvae (*F_L_*) grow in a logistic way without the parasitism pressure from both larval parasitoid (*A*) and facultative pupal hyperparasitoid (*P*). Facultative hyperparasitoid would oviposit in either healthy fly pupae (*F_P_*) or larval primary parasitoid (*A_L_*). See Figure 5a for model diagram and Table 1 for descriptions of all other parameters and their values in the models.

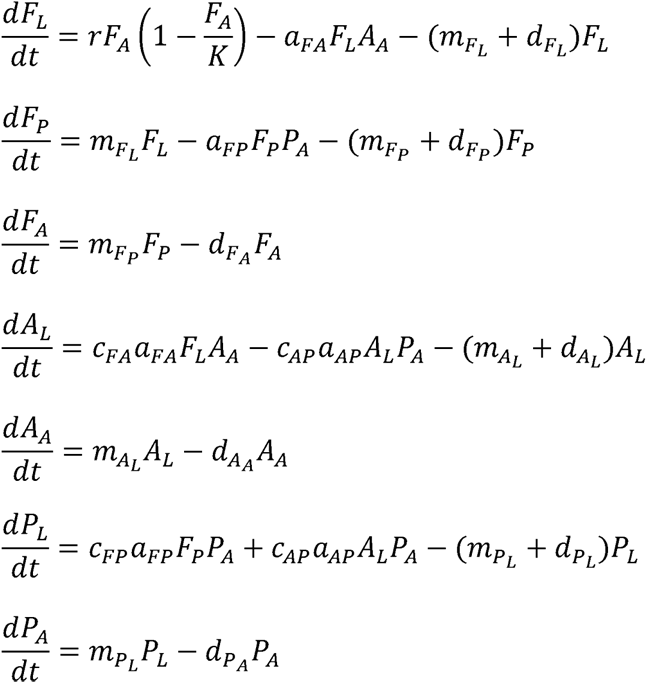

**Table 1.** Parameters and their values in the model.

| Parameters | Description | Values |
| --- | --- | --- |
| $r$ | Intrinsic growth rate of adult fly | 0.1~1.0 in Figure 5b, d<br>1.0 in Figure 5c |
| $K$ | Carrying capacity of adult fly | 50 |
| $m_{FL}$ | Rate of fly larva developing into fly pupa | 1/6 |
| $m_{FP}$ | Rate of fly pupa developing into fly adult | 1/5 |
| $m_{AL}$ | Rate of *A. leverii* larva developing into adult | 1/17 in Figure 5b,<br>1/40~1/3 in Figure 5c, d |
| $m_{PL}$ | Rate of *P. vindemiae* larva developing into adult | 1/21 |
| $a_{ij}$ | Attack rate of species $j$ against species $i$ | $a_{FA} = 0.5$, $a_{FP} = 0.3$<br>$a_{AP} = 0.25$ |
| $c_{ij}$ | Conversion efficiency when species $j$ consuming species $i$ | $c_{FA} = c_{AP} = c_{FP} = c_{AP} = 1$ |
| $d$ | Background mortality | $d_{FL} = d_{FP} = d_{FA} = d_{AL} =$<br>$d_{AA} = d_{PL} = d_{PA} = 0.08$ |

All of the analyses were performed in statistical software R version 4.2.1 (R Core Team 2022).

## 2 Results

### 2.1 Intraguild parasitism promoted species persistence

By comparing *P. vindemiae* exposed to healthy pupae and pupae parasitized by *A. leveri* at larval stage, we found that the number of *P. vindemiae* emerged from pupae with and without *A. leveri* showed no significant difference at low pupae density (GLM: ^2^ = 3.799, *df* = 1, *P* = 0.051, Figure 2a), but significantly more *P. vindemiae* emerged from pupae with *A. leveri* than healthy fly pupae at intermediate (GLM: ^2^ = 541.72, *df* = 1, *P* < 0.001, Figure 2a) or high densities (GLM: ^2^ = 70.93, *df* = 1, *P* < 0.001, Figure 2a). The dissection results showed that at intermediate host density, the number of hosts utilized by *P. vindemiae* with *A. leveri* was greater than that without *A. leveri* (GLM: ^2^ = 53.82, *df* = 1, *P* < 0.001, Figure 2b), and *P. vindemiae* laid more eggs on *A. leveri* than healthy fly pupae when larval flies were exposed to *A. leveri* (GLM: χ = 29.21, *df* = 1, *P* < 0.001, Figure 2b).

**Figure 1.**
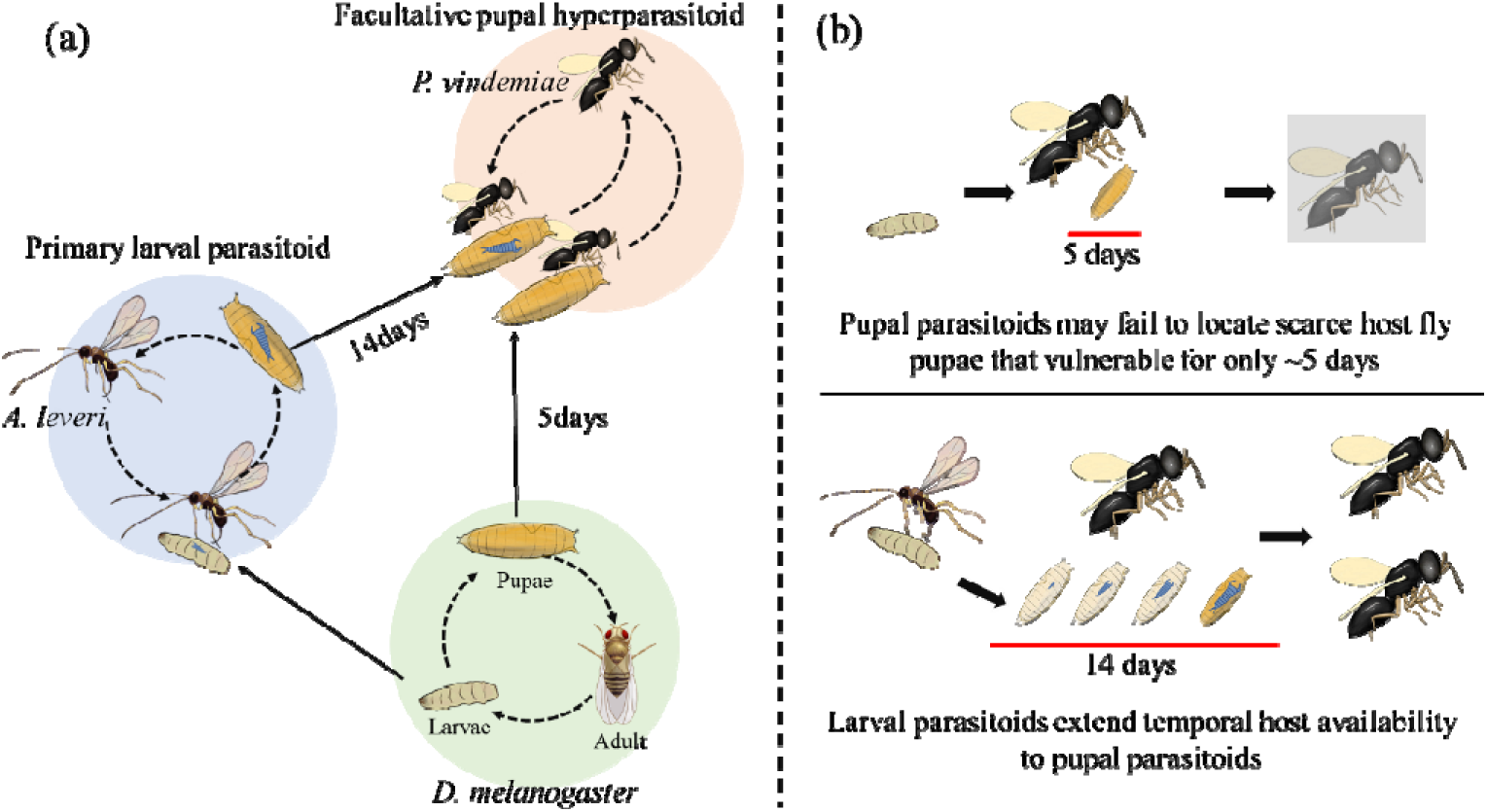
Life history of the studied insects, facultative pupal hyperparasitoid, *Pachycrepoideus vindemiae*, take use of both the healthy *Drosophila* fly pupae and immature larval parasitoids, *Asobara leveri*, within fly pupae as hosts (a). We hypothesize that larval parasitoids promote the persistence of facultative hyperparasitoids by extending temporal host availability (b).

**Figure 2.**
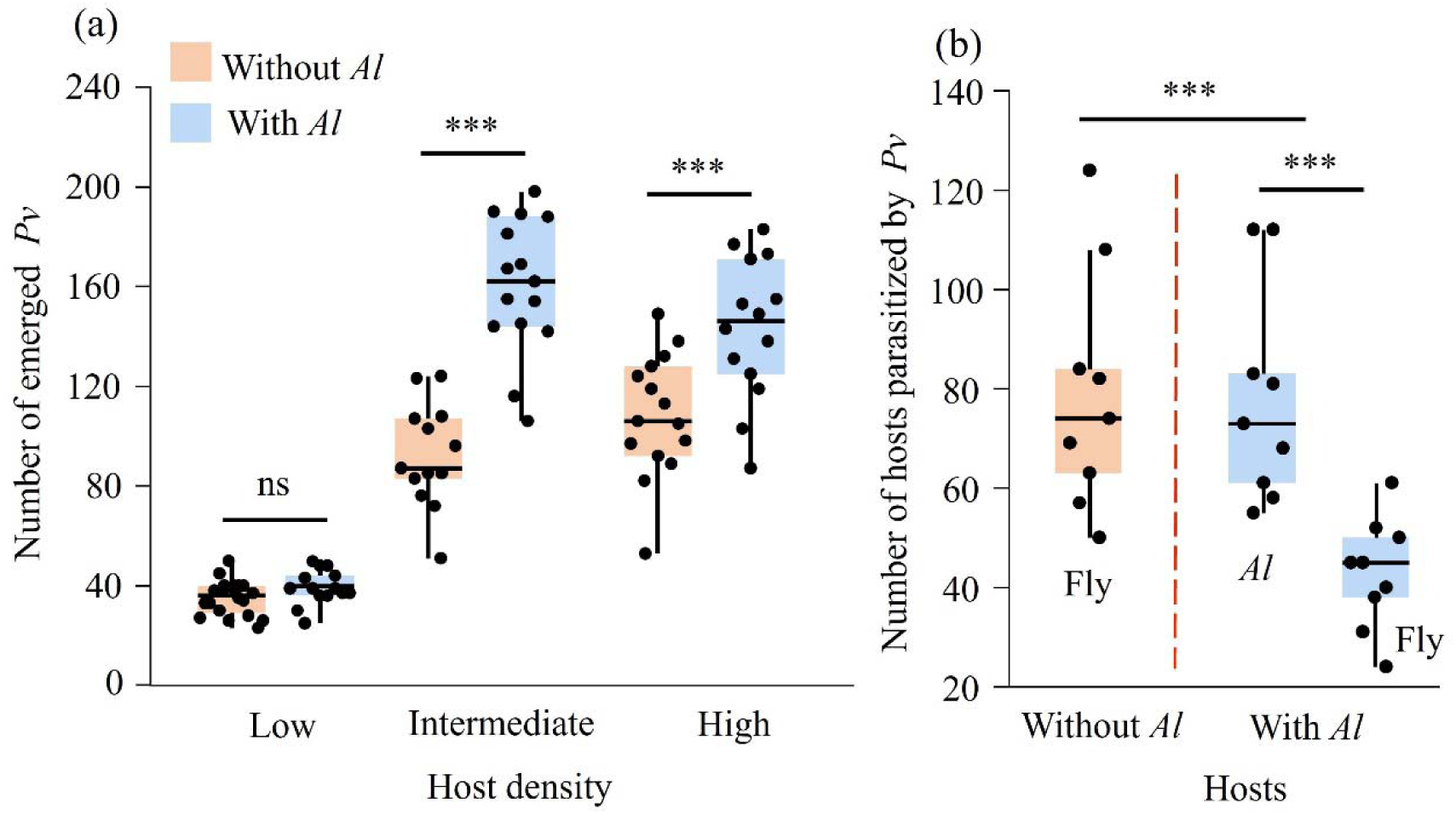
Number of emerged *Pachycrepoideus vindemiae* in different fly pupae density without or with *Asobara leveri*. “***” and “ns” above the boxes indicate significant (*p* < 0.001) and non-significant differences among treatments.

2.2 Effect of *A. leveri* parasitism on the host availability to *P. vindemiae*

The length of immature *A. leveri* within fly pupae that are available to *P. vindemiae* was 16.50 ± 0.20 days (N=30), > 3 times longer than that of healthy *Drosophila* pupae (4.82 ± 0.04 days, N=30, T-tests: *t* = -56.553, *df* = 58, *P* < 0.001, Figure 3a). *P. vindemiae* can successfully emerged as adults from immature *A. leveri* at all development stage, but the survival rate of *P. vindemiae* emerging from 5-day-old *A. leveri* is significantly lower than that from healthy pupae, 10- and 15-day-old *A. leveri* (GLM: χ^2^ = 8.173, *df* = 3, *P* = 0.043, Figure 3b).

**Figure 3.**
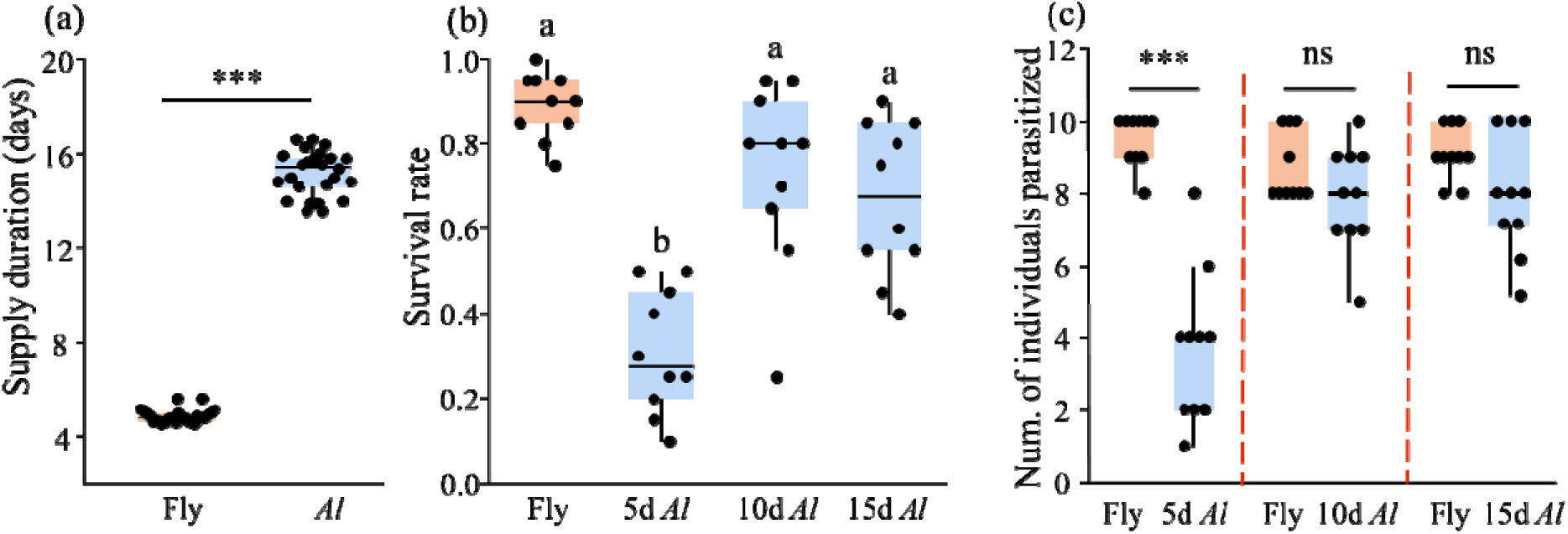
Development time of healthy fly pupae and immature *Asobara leveri* that vulnerable to *Pachycrepoideus vindemiae* (a), survival rate of *P. vindemiae* hosted by immature *A. leveri* of different ages (b), and number of healthy fly pupae and *A. leveri* been parasitized by *Pachycrepoideus vindemiae* in the preference experiment (c). “***” above box indicates significant difference between hosts (*p* < 0.001). Different letters above the boxes indicate significant differences among treatments (*p* < 0.05).

When *P. vindemiae* was exposed to both immature *A. leveri* within fly pupae and healthy fly pupae, *P. vindemiae* preferred to lay eggs in healthy pupae than pupae with 5 days old larval *A. leveri* (GLMM: ^2^ =23.68, *df* = 1, *P* < 0.001, Figure 3c). However, there was no significant difference between the number of healthy pupae and pupae with 10 days old (GLMM: ^2^ = 0.38, *df* = 1, *P* = 0.53) and 15 days old larval *A. leveri* (GLMM: ^2^ = 0.85, *df* = 1, *P* = 0.36).

### 2.3 Fitness of *P. vindemiae* hosted by healthy fly pupae and larval *A. leveri*

*P. vindemiae* offspring hosted by healthy pupae was significantly bigger and have higher fecundity than those hosted by *A. leveri*, especially the old *A. leveri* (Body mass, ANOVA: *F*_2, 33_ = 31.06, *P* < 0.001; Fecundity, GLM: ^2^ = 59.77, *df* = 1, *P* < 0.001, Figure 4a and b).

**Figure 4.**
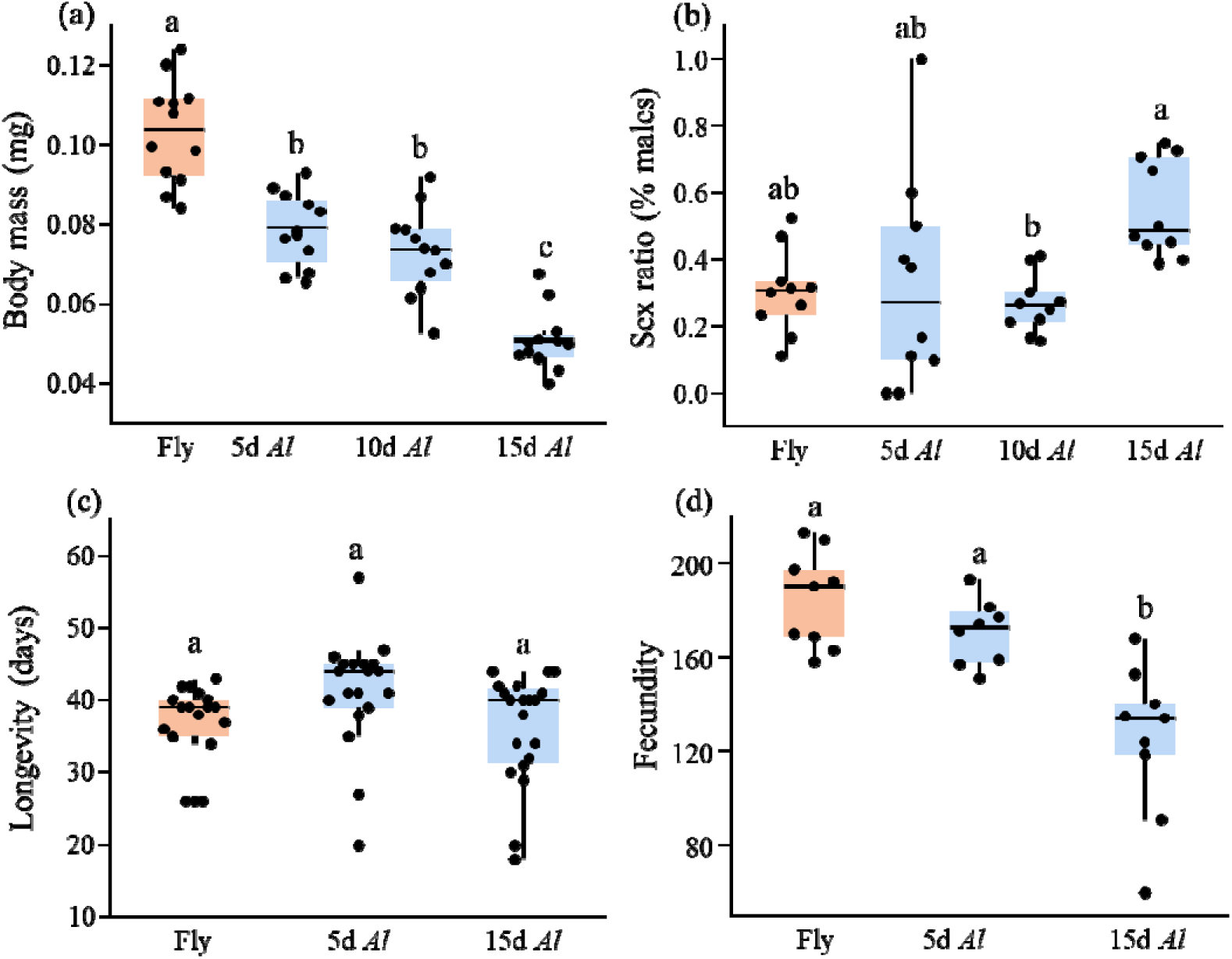
Body mass (a), sex ratio (b), longevity (c) and fecundity (d) of *Pachycrepoideus vindemiae* emerging from healthy fly pupae and *Asobara leveri* of different ages. Different letters above the boxes indicate significant differences among treatments (*p* < 0.05).

In addition, the sex ratio (T-tests: *t* = -1.11, *df* = 38, *P* = 0.276, Figure 4b) and longevity of *P. vindemiae* offspring hosted by *A. leveri* was no significantly different from that of offspring hosted by healthy pupae (GLM: ^2^ = 1.161, *df* = 1, *P* = 0.281, Figure 4c), but sex ratio of *P. vindemiae* offspring increased with the age of *A. leveri* (ANOVA: *F*_2, 27_ = 4.26, *P* = 0.025, Figure 4b).

### 2.4 Model predictions on the coexistence of intraguild parasitism and hyperparasitoids

We then investigated the coexistence of primary parasitoid and hyperparasitoid using theoretical models, where the parasitism-induced extension in the host exposure period was represented by a lower maturing rate of the primary parasitoid (*m_AL_*). In our model, we assumed that the hyperparasitoid had a lower feeding rate on the hosts than primary parasitoid according to observations (Appendix S1: Figure S1). Our modelling results showed facultative hyperparasitoid species cannot survive when fly productivity is low and increase steadily when fly productivity is higher above the threshold, whereas population of primary parasitoids decrease when fly productivity is high enough to support the facultative hyperparasitoids (Figure 5b). In addition, the lower maturing rate of primary parasitoids significantly decreased the minimum fly productivity required for facultative hyperparasitoid to survive (Figure 5c). Consequently, the probability of coexistence of two parasitoid at specific fly productivity level is also depend on the maturing rate of primary parasitoid (Figure 5d).

**Figure 5.**
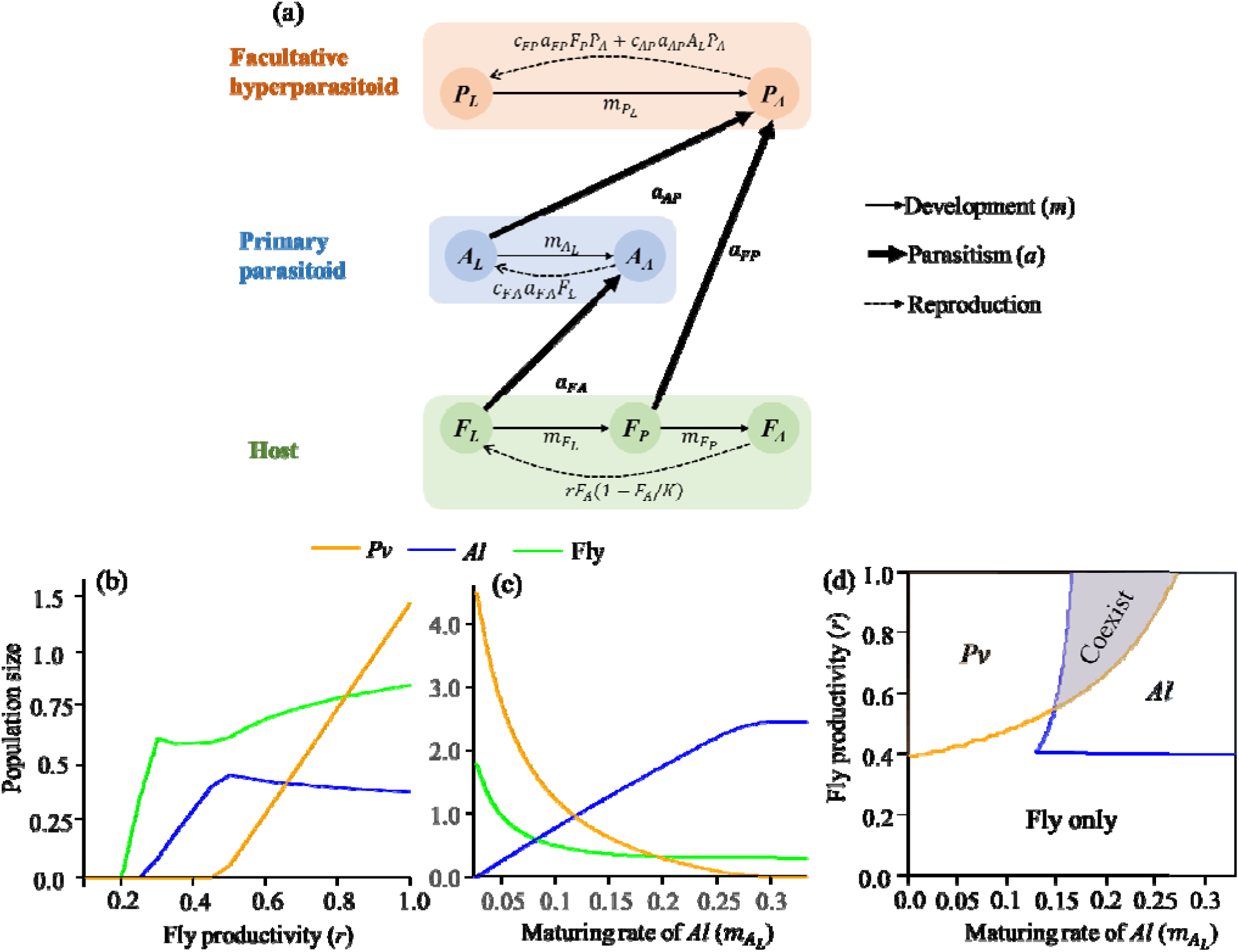
Model diagram used in this study to illustrate parameter settings and the relationships among equations (a). The equilibrium density of host fly, *Asobara leveri* (*Al*) and *Pachycrepoideus vindemiae* (*Pv*) along with host productivity (*r*, b), and the maturing rate of larval parasitoids (*m_AL_*, c), as well as the probability of species persistence with increase of the maturing rate of *A. leveri* (d).

## 3 Discussion

Stage-structure IGP models posits that IG-prey are invulnerable during certain life stages to IG-predators, which is believed to enhance the IGP module persistence by reducing the predation pressure on IG-prey at high resource levels (Borer, 2002). We experimentally revealed that larval parasitoids significantly extend temporal host resource availability to facultative pupal parasitoids, our modeling indicated that intraguild parasitism extended resource availability increased the persistence of pupal parasitoids by decreasing the minimum host density for pupal parasitoids to persist at low host productivity level. These results increased our understanding of how parasitoid diversity is maintained in complex food webs with intraguild parasitism.

In our study, the larval parasitoids significantly extend temporal availability of host resource to pupal parasitoid due to the koinobiont parasitism of *A. leveri* on host pupae (Askari Seyahooei et al., 2020). *Drosophila* fly larvae parasitized by *A. leveri* remain alive till develop into pupae which then become available to pupal parasitoid. Therefore, larval parasitoids do not change the number of fly pupae exposed to *P. vindemiae*. On the other hand, immature *A. leveri* are vulnerable to *P. vindemiae* for ∼14 days during the development within fly pupae, whereas the pupal period of healthy flies that vulnerable to *P. vindemiae* is only ∼5 days. Consequently, *A. leveri* parasitism effectively extend the temporal resource availability to pupal parasitoids. Similar mechanisms may also mediate the species dynamics in the IGP between free-living consumers, as intermediate free-living consumers typically live longer than shared resources (Hutchings, 2021). Thus, although intraguild predation decrease the instant resource availability to top predators (Polis & Myers, 1989), the energy stored in intermediate consumers may become available to top predators in seasons when basal resource is in short supply.

*P. vindemiae* is ectoparasitoid species, adult females lay eggs between the fly pupae shells and fly pupae or immature larval parasitoids, larval *P. vindemiae* absorb nutrients by consuming either developing fly pupae or immature larval parasitoids (Wang & Messing, 2004; Wang et al., 2024). Additionally, although *P. vindemiae* offspring hosted by immature *A. leveri* were smaller in body size and fecundity than those hosted by healthy pupae, adult female pupal parasitoids showed no significant preference for immature primary parasitoids and healthy fly pupae. This could probably because *P. vindemiae* always assesses the host quality based on the size of pupae (Wang et al., 2016), but no significant size variation was observed between healthy pupae and those have been parasitized by *A. leveri*. The fitness reduction of *P. vindemiae* hosted by immature larval parasitoids could be attributed to the loss of nutrient absorbed by fly larvae during transition from *A. leveri* to *P. vindemiae*, as larval parasitoids may induce the melanization immune reaction and resulting in considerable protein loss in host pupae (Yang et al., 2020).

Exploring the mechanisms that determine the length of food chain is one of the central questions in the studies of ecological communities, environmental productivity was thought to be positively correlated with the food chain length (Guo et al., 2023). IGP was traditionally thought to reduce energy flow to consumers at higher trophic levels as some energy is inevitably lost in the metabolism of intermediate consumers (Wang & Brose, 2019), and intensive competition at low productivity may also deplete the resource below the threshold for top predators to survive (Borer & Briggs, 2007). Parasitoid-host food chains are composed by tiny living beings, but up to six trophic levels have been found (Harvey & Wagenaar, 2009). Higher resource use efficiency is believed to provide resource to the consumers in higher trophic levels along the parasitic food chain (Harvey & Wagenaar, 2009; Sanders et al., 2016). Here, our experimental and modeling works clearly demonstrated that the minimum threshold host productivity required for facultative hyperparasitoids to persist decreased linearly with the length of immature period of primary parasitoids. Since most immature primary parasitoids develop for a much longer period than the length of vulnerable period of basal resource, extended temporal resource availability to hyperparasitoids by primary parasitoids should be a widely-relevant mechanisms that increase the persistence of the long parasitic food chains and IGP modules in food webs.

Host developmental stage mediated resource partitioning is considered a mechanism that contributes to the stable coexistence between competing parasitoids without trophic interactions (Godfray, 1994), but parasitoids in the earlier development stage of hosts must be superiority in resource competition for their persistence with facultative hyperparasitoids (Holt & Polis, 1997; Kondoh, 2008; Wang & Brose, 2019). Recent studies demonstrated that dynamic intraguild interactions between two sympatric and congeneric coccinellid species associated with aphids could explain their coexistence in citrus agroecosystems, because IG-prey are thought to dominate the predator community when shared resource is scarce while IG-predators dominate when shared resource is abundant (Bouvet & Urbaneja, 2024). Another recent study revealed that cannibalism among top predators may promote the persistence of top predators at low productivity environment (Bassar et al., 2023; Woodie & Anderson, 2024). Here, we showed that an extended period of host availability to facultative hyperparasitoids also contribute to the network stability at the low resource levels, mainly because *A. leveri* decreased the minimum host flies for *P. vindemiae* to persist. The mechanisms revealed in this study may also elucidate the positive relationship between complexity and stability of parasitoid-host networks where intraguild predation/parasitism modules are commonly existed (Cagnolo et al., 2023).

## Acknowledgments

We thank Tiantian Liu for her help in maintaining the lab insect stocks and Xiaofen Lin for her help in the construction of original IGP model. Mingqiang Wang, Xiaoli Hu and Huijie Qiao provided insightful comments for the earlier manuscript. This study was supported by Fundamental Research Funds for the Central Universities (2024300366) and National Natural Science Foundation of China (32022409).

## Author contributions

XX and HW designed the study. HW conducted all experiments and data analyses. NX, CX, and XX constructed the model, while JL and SW assisted in optimizing it. XX and HW wrote the manuscript, and SW and SS contributed to its revision.

## References

Amarasekare, P. 2007. Trade-offs, temporal variation, and species coexistence in communities with intraguild predation. Ecology 88: 2720–2728.

Arim, M., and P. A. Marquet. 2004. Intraguild predation: a widespread interaction related to species biology. Ecology Letters 7: 557–564.

Askari Seyahooei, M., K. Kraaijeveld, A. Bagheri, and J. J. M. van Alphen. 2020. Adult size and timing of reproduction in five species of Asobara parasitoid wasps. Insect Science 27: 1334–1345.

Banerji, A., and P. J. Morin. 2014. Trait-mediated apparent competition in an intraguild predator–prey system. Oikos 123: 567–574.

Bassar, R. D., T. Coulson, and J. Travis. 2023. Size-dependent intraguild predation, cannibalism, and resource allocation determine the outcome of species coexistence. The American Naturalist 201: 712–724.

Borer, E. T. 2002. Intraguild predation in larval parasitoids: implications for coexistence. Journal of Animal Ecology 71: 957–965.

Borer, E. T., C. J. Briggs, and R. D. Holt. 2007. Predators, parasitoids, and pathogens: a cross-cutting examination of intraguild predation theory. Ecology 88: 2681–2688.

Borer, E. T., C. J. Briggs, W. W. Murdoch, and S. L. Swarbrick. 2003. Testing intraguild predation theory in a field system: does numerical dominance shift along a gradient of productivity? Ecology Letters 6: 929–935.

Bouvet, J. P. R., A. Urbaneja, and C. Monzo. 2024. Dynamic intraguild interactions between two sympatric and congeneric coccinellid species associated with aphids could explain their coexistence in citrus agroecosystems. Biological Control 192: 105506.

Cagnolo, L., L. Bernaschini, A. Salvo, and G. Valladares. 2023. Habitat area and edges affect the length of trophic chains in a fragmented forest. Journal of Animal Ecology 92: 2067–2077.

Chesson, P., and J. J. Kuang. 2008. The interaction between predation and competition. Nature 456: 235–238.

Flick, A. J., T. A. Coudron, and B. D. Elderd. 2020. Intraguild predation decreases predator fitness with potentially varying effects on pathogen transmission in a herbivore host. Oecologia 193: 789–799.

Frago, E. 2016. Interactions between parasitoids and higher order natural enemies: intraguild predation and hyperparasitoids. Current Opinion in Insect Science 14: 81–86.

Godfray, H. C. J. 1994. Parasitoids: behavioral and evolutionary ecology. Princeton University Press, Princeton, New Jersey, USA.

Guo, G. M., G. Barabás, G. Takimoto, D. Bearup, W. F. Fagan, D. D. Chen, and J. B. Liao. 2023. Towards a mechanistic understanding of variation in aquatic food chain length. Ecology Letters 26: 1926–1939.

Gutgesell, M. K., K. S. McCann, G. Gellner, K. Cazelles, C. J. Greyson-Gaito, C. Bieg, M. M. Guzzo, et al., 2022. On the dynamic nature of omnivory in a changing world. Bioscience 72: 416–430.

Harvey, J. A., R. Wagenaar, and T. M. Bezemer. 2009. Interactions to the fifth trophic level: secondary and tertiary parasitoid wasps show extraordinary efficiency in utilizing host resources. Journal of Animal Ecology 78: 686–692.

Hassell, M. P. 2000. The spatial and temporal dynamics of host-parasitoid interactions. Oxford: Oxford University Press.

Holling, C. S. 1959. Some characteristics of simple types of predation and parasitism1. The canadian entomologist 91: 385–398.

Holt, R. D., and G. A. Polis. 1997. A theoretical framework for intraguild predation. The American Naturalist 149: 745–764.

Hutchings, J. A. 2021. A Primer for Life-Histories: Ecology, Evolution, and Applications. Oxford: Oxford University Press.

Jervis, M. A., and N. A. C. Kidd. 1986. Host-feeding strategies in hymenopteran parasitoids. Biological Reviews 61: 395–434.

Julious, S. A. 2004. Using confidence intervals around individual means to assess statistical significance between two means. Pharmaceutical Statistics 3: 217–222.

Kondoh, M. 2008. Building trophic modules into a persistent food web. Proceedings of the National Academy of Sciences 105: 16631–16635.

Kratina, P., R. M. LeCraw, T. Ingram, and B. R. Anholt. 2012. Stability and persistence of food webs with omnivory: Is there a general pattern? Ecosphere 3: 1–18.

Lafferty, K. D., S. Allesina, M. Arim, C. J. Briggs, G. De Leo, A. P. Dobson, J. A. Dunne, et al., 2008. Parasites in food webs: the ultimate missing links. Ecology Letters 11: 533–546.

Liao, J., D. Bearup, and W. F. Fagan. 2020. The role of omnivory in mediating metacommunity robustness to habitat destruction. Ecology 101: e03026.

Liao, J., D. Bearup, Y. Wang, I. Nijs, D. Bonte, Y. Li, U. Brose, et al., 2017. Robustness of metacommunities with omnivory to habitat destruction: disentangling patch fragmentation from patch loss. Ecology 98: 1631–1639.

Lucas, É., D. Coderre, and J. Brodeur. 1998. Ingraguild predation among aphid predators: Characterization and influence of extraguild prey density. Ecology 79: 1084–1092.

Lue, C. H., M. L. Buffington, S. Scheffer, M. Lewis, T. A. Elliott, A. R. I. Lindsey, A. Driskell, et al., 2021. DROP: Molecular voucher database for identification of *Drosophila* parasitoids. Molecular Ecology Resourources 21: 2437–2454.

Mariano-Macedo, A., Y. M. Vázquez-González, A. M. Martinez, A. Rebollar-Alviter, J. I. Figueroa, S. I. Morales, E. Vinuela, et al., 2020. Biological traits of a Pachycrepoideus vindemiae Mexican population on the host *Drosophila* suzukii. Bulletin of Insectology 73.

May, R. M. 1977. Predators that switch. Nature 269: 103–104.

McCann, K., and A. Hastings. 1997. Re–evaluating the omnivory–stability relationship in food webs. Proceedings of the Royal Society of London. Series B: Biological Sciences 264: 1249–1254.

McMeans, B. C., K. S. McCann, M. Humphries, N. Rooney, and A. T. Fisk. 2015. Food Web Structure in Temporally-Forced Ecosystems. Trends in ecology & evolution 30: 662–672.

Moeller, H. V., M. G. Neubert, and M. D. Johnson. 2019. Intraguild predation enables coexistence of competing phytoplankton in a well mixed water column. Ecology 100: e02874.

Morin, P. 1999. Productivity, intraguild predation, and population dynamics in experimental food webs. Ecology 80: 752–760.

Müller, C. B., and J. Brodeur. 2002. Intraguild predation in biological control and conservation biology. Biological Control 25: 216–223.

Murdoch, W. W., and A. Oaten. 1975. Predation and Population Stability. In Advances in ecological research, Edited by A. MacFadyen, 1–131. USA: Academic Press.

Navarrete, S. A., B. A. Menge, and B. A. Daley. 2000. Species Interactions in Intertidal Food Webs: Prey or Predation Regulation of Intermediate Predators? Ecology 81: 2264–2277.

Polis, G. A., C. A. Myers, and R. D. Holt. 1989. The ecology and evolution of intraguild predation: Potential competitors that eat each other. Annual Review of Ecology and Systematics 20: 297–330.

Rayl, N. D., G. Bastille-Rousseau, J. F. Organ, M. A. Mumma, S. P. Mahoney, C. E. Soulliere, K. P. Lewis, et al., 2018. Spatiotemporal heterogeneity in prey abundance and vulnerability shapes the foraging tactics of an omnivore. Journal of Animal Ecology 87: 874–887.

Rosenheim, J. A. 2007. Intraguild predation: New theoretical and empirical perspectives. Ecology 88: 2679–2680.

Rosenheim, J. A., H. K. Kaya, L. E. Ehler, J. J. Marois, and B. A. Jaffee. 1995. Intraguild predation among biological-control agents: Theory and evidence. Biological Control 5: 303–335.

Sanders, D., A. Moser, J. Newton, and F. J. F. van Veen. 2016. Trophic assimilation efficiency markedly increases at higher trophic levels in four-level host–parasitoid food chain. Proceedings of the Royal Society of London B: Biological Sciences 283: 20153043.

Sentis, A., J. L. Hemptinne, J. Brodeur, and M. Eubanks. 2014. Towards a mechanistic understanding of temperature and enrichment effects on species interaction strength, omnivory and food web structure. Ecology Letters 17: 785–793.

Sieber, M., and F. M. Hilker. 2011. Prey, predators, parasites: intraguild predation or simpler community modules in disguise? Journal of Animal Ecology 80: 414–421.

Thompson, C. M., and E. M. Gese. 2007. Food webs and intraguild predation: community interactions of a native mesocarnivore. Ecology 88: 334–346.

Vance-Chalcraft, H. D., J. A. Rosenheim, J. R. Vonesh, C. W. Osenberg, and A. Sih. 2007. The influence of intraguild predation on prey suppression and prey release: A meta-analysis. Ecology 88: 2689–2696.

Wang, H., T. Liu, S. Sun, O. T. Lewis, and X. Xi. 2024. Temporal variability in host availability alters the outcome of competition between two parasitoid species. Journal of Animal Ecology 93: 1845–1853.

Wang, S., U. Brose, and D. Gravel. 2019. Intraguild predation enhances biodiversity and functioning in complex food webs. Ecology 100: e02616.

Wang, X., B. N. Hogg, A. Biondi, and K. M. Daane. 2021. Plasticity of body growth and development in two cosmopolitan pupal parasitoids. Biological Control 163: 104738.

Wang, X., G. Kaçar, A. Biondi, and K. M. Daane. 2016. Foraging efficiency and outcomes of interactions of two pupal parasitoids attacking the invasive spotted wing *Drosophila*. Biological Control 96: 64–71.

Wang, X., and R. H. Messing. 2004. The ectoparasitic pupal parasitoid, Pachycrepoideus vindemmiae (Hymenoptera: Pteromalidae), attacks other primary tephritid fruit fly parasitoids: host expansion and potential non-target impact. Biological Control 31: 227–236.

Woodie, C. A., and K. E. Anderson. 2024. Preferential cannibalism as a key stabilizing mechanism of intraguild predation systems with trophic polymorphic predators. Theoretical Ecology 17: 59–72.

Xi, X., N. Eisenhauer, and S. Sun. 2015. Parasitoid wasps indirectly suppress seed production by stimulating consumption rates of their seed-feeding hosts. Journal of Animal Ecology 84: 1103–1111.

Yang, L., L. M. Qiu, Q. Fang, D. W. Stanley, and G. Y. Ye. 2020. Cellular and humoral immune interactions between *Drosophila* and its parasitoids. Insect Science 28: 1208–1227.

Zheng, J., U. Brose, D. Gravel, B. Gauzens, M. Luo, and S. Wang. 2021. Asymmetric foraging lowers the trophic level and omnivory in natural food webs. Journal of Animal Ecology 90: 1444–1454.

