## Supplemental file for "Intraguild parasitism promotes the persistence of facultative hyperparasitoids by extending temporal host availability"

**Abstract:**

Intraguild predation (IGP) is pervasive in natural food webs, although theoretical models predict restricted parameter space for the coexistence of IG-prey and IG-predators. One potential mechanism is that IGP persistence might be increased when IG-prey extend resource availability for IG-predators. Here we test this hypothesis using manipulative experiments in combination with mathematical modeling. Our experiment system includes *Drosophila* flies (*Drosophila melanogaster*, basal resources), primary larval parasitoids (*Asobara leveri*, IG-prey) and facultative pupal hyperparasitoid (*Pachycrepoideus vindemiae*, IG-predator). We measured the performance of facultative pupal parasitoids with and without the primary larval parasitoid. Our study revealed that larval parasitoids significantly increased the number of pupal parasitoids emerged from limited host resources, due to the extended resource availability to facultative hyperparasitoid. Specifically, immature primary parasitoids within *Drosophila* pupae can be explored by facultative hyperparasitoids for up to 14 days, while the healthy fly pupae are available to hyperparasitoids for only 5 days. Despite a significant reduction in offspring fitness of the facultative hyperparasitoids when reared on larval parasitoids versus healthy *Drosophila* pupae, adult female facultative hyperparasitoids showed no significant preference for healthy fly pupae over immature primary parasitoids. Our theoretical models demonstrated that an extended development period of the primary parasitoid can promote the persistence of the facultative hyperparasitoids and allow their coexistence with the primary parasitoid, although too prolonged development could drive the extinction of the primary parasitoid. As IG-prey often take longer time to develop after consuming the shared resources, extension of temporal resource availability could be a general mechanism contribute to the persistence of intraguild predators.

**Key words:** Predation and competition, *Drosophila* fly, Stage-structure, Biological control, Ontogeny niche

**1 Materials and methods**

**1.1 host use efficiency of larval and pupal parasitoids**

To estimate the host-searching efficiency of *A. leveri*, we investigated the functional response of *A. leveri* to varying host larval densities (20, 40, 60, 80, 100). These host densities were ascertained through preliminary tests to ensure that parasitoids had access to enough hosts for a specified period. We released 1, 2, 3, 4, and 5 newly emerged and mated female *D. melanogaster* into vials to lay eggs for 24 hours. After the eggs developed into 2-day-old larvae, we introduced one pair of *A. leveri* into the vial to oviposit for 24hours. Once the larvae pupated, the number of unparasitized and parasitized pupae was then recorded. The puparia parasitized by *A. leveri* were darker than those of healthy ones.

To estimate the host-searching efficiency of *P. vindemiae*, we investigated the functional response of *P. vindemiae* to varying host pupal density (3, 6, 9, 12, 15, 18). To ensure enough pupae, we released 4 mated *D. melanogaster* females into the vial to lay eggs for 24 hours. Once the larvae had pupated, the fresh pupae were transferred to vials with expected host densities. A pair of *P. vindemiae* was then introduced into the vial to oviposit for 24 hours, they were fed 10% sucrose water solution. 10 days after parasitism, unparasitized and parasitized pupae were counted. At this stage, the larvae of *P. vindemiae* in the parasitized pupae could be easily observed. A total of 7 replicates were conducted for each host density treatment for both *A. leveri* and *P. vindemiae*.

The number of hosts parasitized by two parasitoid species increased with host density and then remains constant, thus we employed the type II functional response function (Holling 1959) to estimate the searching efficiency (*a*) and handling time (*T_h_*) of both *P. vindemiae* and *A. leveri* as follows:

$$N_{a}={(aT_{t}P_{t}N}_{0}) / [1+aT_{h}N_{0}]$$

where *Pt* is the number of parasitoids (=1), and *T_t_* is the total time (= 1d) available for parasitoids to exploit hosts, *N_a_* is the number of parasitized hosts, and *N_0_* is the number of hosts. Statistical significant difference was revealed when 84% confidence intervals of *a* between two parasitoid species do not overlap (Julious 2004).

**1.2 Statistical analyses**

We employed generalized linear models (GLMs) with Poisson error distribution (link = “log”) to test if the number of *P. vindemiae* and *A. leveri* significantly increase with host density. A one-way analysis of variance (ANOVA) was used to examine if the fitting function of *P. vindemiae* and *A*. *leveri* better than the function y = 0.

All statistical analyses were processed using the statistical software R version 4.2.1.

**2 Results**

The number of hosts parasitized by both *P. vindemiae* and *A. leveri* per day increased with host density (GLM: *P. vindemiae*: *χ*^2^ = 70.10, *df* = 1, *P* < 0.001; *A. leveri*: *χ*^2^ = 180.49, *df* = 1, *P* < 0.001, Figure S1), before host density reaches 18 and 130, respectively. The functional response equation of *P. vindemiae* parasitizing fly pupae was *Na* = 0.83*N_0_* / (1 + 0.019*N_0_*) (ANOVA: *F*_2, 4_ = 208.40, *P* < 0.001, *R*^2^ = 0.62). The functional response equation of *A. leveri* parasitizing fly larvae was *Na* = 1.31*N_0_* / (1 + 0.007*N_0_*) (ANOVA: *F*_2, 35_ = 900.83, *P* < 0.001, *R*^2^ = 0.90). The searching efficiency (*a*) of *P. vindemiae* (0.83 ± 0.18, 84% C.I: 0.79 ~ 0.88) is significantly lower than *A. leveri* (1.31 ± 0.14, 84% C.I.: 1.28 ~ 1.35).

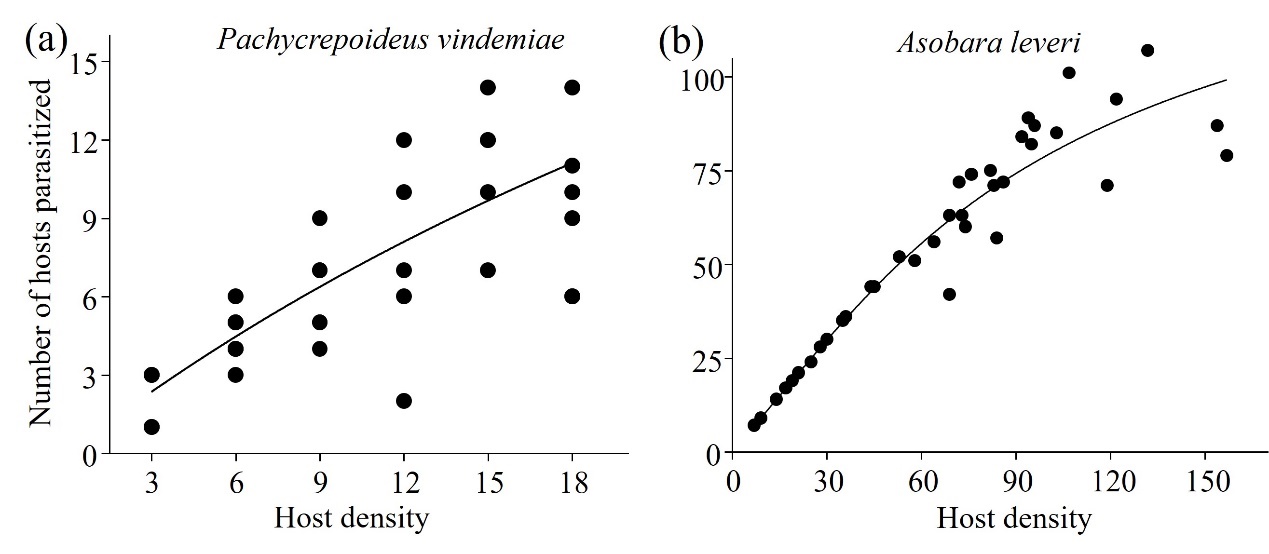

Figure S1 Type II functional response of *Pachycrepoideus vindemiae* (a) and *Asobara leveri* (b) parasitizing *Drosophila* host.

**References：**

Holling, C. S. 1959. Some characteristics of simple types of predation and parasitism1. *The canadian entomologist* 91: 385-398.

Julious, S. A. 2004. Using confidence intervals around individual means to assess statistical significance between two means. *Pharmaceutical Statistics* 3: 217-222.
